# Cognitive load modulates cortical excitability in the human prefrontal cortex

**DOI:** 10.64898/2026.04.25.720856

**Authors:** Umair Hassan, Lily Forman, James W. Hartford, Hansong Lee, Sara Parmigiani, Nai-Feng Chen, Jessica M. Ross, Ethan A. Solomon, Juha Gogulski, Naryeong Kim, Jade Truong, Christopher C. Cline, Corey J. Keller

## Abstract

The prefrontal cortex dynamically adjusts its excitability to meet changing cognitive demands, yet our understanding of how these rapid state changes unfold in real time remains limited. Existing evidence relies predominantly on correlational neuroimaging and invasive electrophysiology in clinical populations, neither of which affords causal measurement in healthy humans. Here, using concurrent single-pulse TMS-EEG during a parametric cognitive multi-source interference task (MSIT) in 27 healthy participants, we causally probed left prefrontal excitability across conditions spanning a range of cognitive interference levels. Prefrontal excitability scaled systematically with cognitive load, peaking at intermediate incongruency levels where trial-type uncertainty was greatest, and differentiating high-from low-conflict trials at the single-subject level. Critically, prefrontal excitability and midfrontal theta power showed parallel state dependency patterns, linking causal perturbation of prefrontal circuit responsiveness to an established oscillatory marker of cognitive control. The observed excitability modulation was selective to the left prefrontal cortex: stimulation of right dlPFC, pre-SMA, and vertex in a separate control cohort (N=5) produced no load-dependent effects. These results position prefrontal excitability as a real-time, regionally selective physiological readout of how the human prefrontal cortex adapts to cognitive demand.

**Highlights:**

- We evaluated the links between cognitive interference, prefrontal excitability, and intrinsic theta activity during a multi-source interference task
- Cognitive load, estimated based on behavioral performance, was greatest at at intermediate interference levels (33% congruent), where trial-type uncertainty was maximal
- Prefrontal excitability, measured via single-pulse TMS-evoked potentials, scaled with cognitive load
- Effects were observed at the single subject level and were specific to prefrontal stimulation
- Prefrontal excitability changes and midfrontal theta power showed parallel scaling with cognitive load, linking causal probing to established oscillatory markers
- This noninvasive technique provides causal measurement of prefrontal engagement during cognitive processing
- The methodology demonstrates potential for tracking brain state dynamics in both healthy cognition and neuropsychiatric disorders

## 1. Introduction

The prefrontal cortex dynamically adjusts neural activity to meet changing cognitive demands, yet how these rapid state changes unfold remains inaccessible to current noninvasive methods.. This poses a fundamental barrier in cognitive neuroscience and clinical translation alike. Prefrontal dysfunction underlies major psychiatric disorders including depression, schizophrenia, and ADHD (Arnsten, 2011; Gratton et al., 2024), and brain stimulation therapies increasingly target prefrontal circuits (Keller et al., 2018; Gogulski et al., 2024a), yet we cannot currently verify whether these interventions engage their intended circuits during active cognition. Functional neuroimaging reveals the spatial architecture of prefrontal control networks (Gratton et al., 2024) and electrophysiology identifies oscillatory signatures such as midfrontal theta (Cavanagh & Frank, 2014; Cavanagh & Shackman, 2015), but neither approach can causally probe how cortical excitability shifts as cognitive demands change. Intracranial recordings offer causal precision but are limited to rare clinical populations (Smith et al., 2019; Wang et al., 2024). A noninvasive method that can track prefrontal excitability changes with millisecond precision during cognitive processing would advance both basic understanding of cognitive control and the development of state-informed therapeutic interventions.

Transcranial magnetic stimulation combined with electroencephalography (TMS-EEG) addresses these limitations by delivering causal perturbations while recording cortical responses with millisecond resolution. Single TMS probe pulses evoke early-local TMS-evoked potentials (EL-TEPs), short-latency responses within 20-80 ms that index local cortical excitability at the stimulation site (Keller et al., 2018; Belardinelli et al., 2021). Recent work has established these EL-TEPs as robust and reliable markers of prefrontal excitability (Gogulski et al., 2024a; Gogulski et al., 2024b; Parmigiani et al., 2025), with responses that inversely relate to non-neural artifacts, supporting their neurophysiological origin (Gogulski et al., 2024a). However, whether EL-TEPs can track the rapid excitability changes that accompany cognitive state transitions remains untested. To address this, we leveraged the multi-source interference task (MSIT), which parametrically manipulates cognitive interference through varying levels of response conflict (Bush & Shin, 2006). Critically, interference tasks engage midfrontal theta oscillations (Cavanagh & Frank, 2014; Cavanagh & Shackman, 2015), providing an established neural benchmark against which to evaluate EL-TEP sensitivity to cognitive state. Furthermore, prior studies established EL-TEP reliability by systematically varying TMS input parameters (e.g., intensity, coil orientation), leaving open the question of whether TMS-evoked potentials are sensitive to endogenous changes in cortical state (Conde et al., 2019). The present investigation addresses this gap: by holding all TMS input parameters constant while experimentally modulating prefrontal cognitive state, any resulting EL-TEP variability can be attributed to changes in underlying cortical excitability rather than differences in the stimulation itself, providing a stronger test of EL-TEPs’ physiological state-dependency.

That prefrontal excitability adapts to cognitive demands has long been inferred from correlational neuroimaging and invasive recordings in clinical populations, but direct causal evidence in healthy humans has been lacking. Here, we provide noninvasive causal evidence that left prefrontal excitability selectively tracks cognitive load during active cognitive processing. Using concurrent single-pulse TMS-EEG during the MSIT in 27 healthy participants, we measured prefrontal excitability across parametrically varied cognitive interference conditions. Prefrontal excitability scaled systematically with cognitive load, peaking at intermediate incongruency levels where trial-type uncertainty was greatest. This modulation was selective to the left prefrontal cortex: stimulation of right dlPFC, pre-SMA, and vertex in a separate control cohort (N=5) produced no load-dependent effects. Importantly, prefrontal excitability and midfrontal theta power showed parallel state dependent patterns, linking causal measurement of circuit responsiveness to an established oscillatory marker of cognitive control. These findings demonstrate that the left prefrontal excitability is dynamically and selectively regulated by cognitive demands, establishing a real-time causal window into how the human prefrontal cortex adapts to changing task conditions.

## 2. Methods

### 2.1. Participants

Forty-eight healthy participants (19-63 years old, mean = 40.21, SD = 12.87, 24 female) initially provided written informed consent under a protocol approved by the Stanford University Institutional Review Board. Inclusion criteria required participants to be aged 18-65 years, fluent in English, able to travel to the study site, and fully vaccinated against COVID-19. All participants completed the Quick Inventory of Depressive Symptomatology (16-item, QIDS) self-report questionnaire, with exclusion for scores of 11 or higher indicating moderate or more severe depression. Additional exclusion criteria included lifetime history of psychiatric or neurological disorder, substance abuse or dependence in the past month, recent heart attack (< 3 months), pregnancy, presence of absolute contraindications for repetitive TMS or MRI, and history of psychotropic medication use following established TMS safety guidelines (Rossi et al., 2021). Eight participants were excluded during initial consent based on these criteria, and two were lost to follow-up. Additional participants were excluded due to excessive EEG artifacts or incomplete data collection. The final sample for the main experiment consisted of 27 participants who completed all 11 cognitive interference conditions with concurrent left dlPFC stimulation across both study visits (Table S1). A separate control experiment was conducted in 5 additional participants who received TMS at four cortical sites (left dlPFC, right dlPFC, left pre-SMA, and vertex) across three cognitive interference conditions (MSIT control, 100% congruent, and 33% congruent). All participants provided written informed consent and received financial compensation for their participation.

### 2.2. Multi-Source Interference Task (MSIT)

The MSIT is a well-validated cognitive-interference paradigm that combines elements of the Stroop (Stroop, 1935), Flanker (Eriksen & Eriksen, 1974), and Simon (Simon, 1990) tasks to systematically manipulate cognitive-control demands as originally developed and validated by Bush and colleagues (Bush et al., 2003; Bush & Shin, 2006). On each trial, participants viewed three numbers displayed horizontally on a computer screen and were instructed to identify the value of the unique number (the number that differed from the other two) by pressing the corresponding button on a response pad using their right hand (index = position 1, middle = 2, ring = 3) (Figure 1A–B).

**Figure 1:**
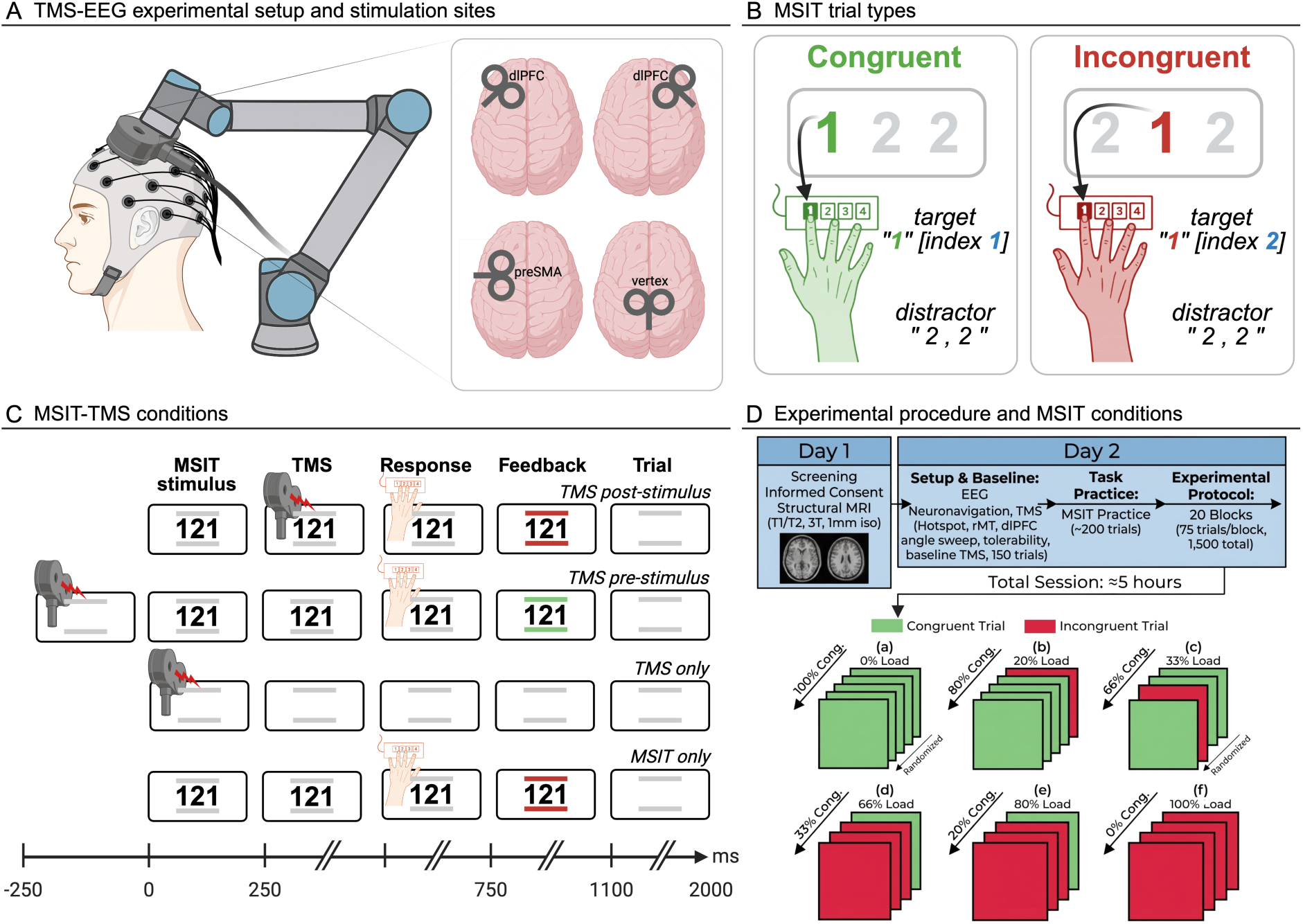
Experimental design and cognitive interference manipulation during concurrent TMS-EEG. (A) Schematic of the experimental setup showing concurrent transcranial magnetic stimulation (TMS) and electroencephalography (EEG) during the Multi-Source Interference Task (MSIT). Four brain regions are stimulated: left dorsolateral prefrontal cortex (ldlPFC), right dlPFC (rdlPFC), vertex, and left pre-supplementary motor area (pre-SMA). The setup includes the participant wearing an EEG cap, the TMS coil positioned by a robotic arm, and the task display. (B) MSIT trial structure and timing. Participants viewed three numbers and identified the unique number’s position by button press. Examples show congruent trials (green, e.g., "1 2 2" with target "1" at index 1) where number identity matches position, and incongruent trials (red, e.g., "2 1 2" with target "1" at index 2) where identity conflicts with position, creating multi-source interference. (C) Timeline of different MSIT trial types with respect to when TMS is delivered. The different rows illustrate variations in trial structure: TMS delivered after the stimulus, TMS delivered before the stimulus, TMS alone with no task (no visual stimulus or response), and task alone with no TMS. (D) Study timeline across two days. Day 1: Screening, informed consent, structural MRI (T1/T2, 3T scanner, 1 mm isotropic resolution). Day 2: Setup included EEG cap placement, baseline TMS measurement (neuronavigation, TMS hotspot identification, resting motor threshold determination, dlPFC target localization, single sweep tolerability check, baseline TMS with 150 trials), followed by MSIT practice (∼200 trials). Experimental protocol consisted of 20 blocks (75 trials/block, 1,500 total trials) with approximately 5-hour total session duration. Cognitive interference was systematically varied across blocks by manipulating the ratio of congruent to incongruent trials from 100% congruency conditions to 0% congruence condition blocks. Green frames represent congruent trials; red frames represent incongruent trials. Block order was pseudorandomized across participants, with brief rest breaks between blocks.

In congruent trials, the identity of the unique number matched its spatial position on the screen and response button (e.g., "1-0-0" requires a left-button press for position 1). In incongruent trials, the position of the unique number conflicted with its identity (e.g., "2-1-2" requires a left-button press because the unique item is “1”, but position 1 contains "2"). This design creates multi-source interference by combining stimulus-stimulus, stimulus-response, and spatial conflict. The MSIT has been validated through fMRI studies, which demonstrate that it reliably increases BOLD response in prefrontal networks, with BOLD response scaling with cognitive load (Bush & Shin, 2006). This task robustly engages the cognitive control network, producing reliable behavioral interference effects and consistent BOLD fMRI activation of dorsal anterior cingulate and dorsolateral prefrontal cortex. The MSIT has been shown to be altered in depression and other psychiatric disorders (Bush et al., 2008). Importantly, the task allows parametric manipulation of cognitive interference while keeping sensory inputs relatively constant, providing an ideal testbed for examining whether noninvasive measures of prefrontal excitability tracks cognitive state changes. By combining this behavioral paradigm with precisely-timed TMS pulses, we can directly test whether causal probing captures the dynamic engagement of prefrontal circuits during cognitive control. Furthermore, given that interference tasks may reliably engage prefrontal theta oscillations this paradigm allows us to investigate the relationship between TMS-elicited evoked neural responses and intrinsic rhythms that coordinate prefrontal function.

Task parameters were as follows: fixation = 500 ms, stimulus = 1,000 ms, feedback = 250 ms, inter-trial interval = 2,250 ms (Figure 1C). Single-pulse TMS was delivered 250 ms after stimulus onset, timed to occur during cognitive processing but before motor execution (Taylor et al., 2008). Trials that had a response before this time were rejected (1.6% of trials). Cognitive interference was manipulated block-wise by varying the proportion of congruent to incongruent trials, allowing expectation-based modulation of control engagement.

### 2.3. Study Design and Experimental Protocol

In the main experiment, all 27 participants completed two study visits on separate days. Prior to the experimental session, participants underwent screening procedures, provided informed consent, and completed T1 and T2 weighted structural MRI scans (3 T scanner, 1 mm isotropic resolution) for subsequent TMS neuronavigation. The experimental session (Day 2) began with EEG cap placement and TMS targeting procedures, including motor hotspot identification, resting motor threshold (rMT) determination, dlPFC target localization, and coil angle optimization to minimize muscle artifacts. Following these setup procedures, participants completed a baseline resting-state block consisting of 150 single-pulse TMS trials delivered to the left dorsolateral prefrontal cortex while at rest with eyes open (Figure 1D).

Participants then received verbal and written instructions for the Multi-Source Interference Task (MSIT) and completed a brief practice block (≈ 20 trials) to familiarize themselves with the task requirements and button-response mappings. The main experimental protocol consisted of 20 blocks, each containing 75 trials (1,500 total), with cognitive interference systematically varied across blocks by manipulating the ratio of congruent to incongruent trials (Bush & Shin, 2006; Bush et al., 2003). Block types included: 100% congruent (0% incongruent), 80% congruent (20% incongruent), 66% congruent (33% incongruent), 33% congruent (66% incongruent), 20% congruent (80% incongruent), and 0% congruent (100% incongruent). Block order was pseudorandomized across participants to control for order effects and fatigue. Two control blocks (5th and 11th) featured a 0-back working-memory task rather than the MSIT, and two additional blocks (10th and 20th) presented MSIT trials without TMS to assess behavioral performance without concurrent brain stimulation. Brief rest breaks (10-20 minutes) were provided between blocks to maintain participant comfort, vigilance, and attention throughout the approximately 5-hour experimental session. In the separate control experiment (N=5), participants underwent an identical setup and received single-pulse TMS at four cortical sites (left dlPFC, right dlPFC, left pre-SMA, and vertex) during three MSIT conditions (control, 100% congruent, and 33% congruent).

### 2.4. Transcranial Magnetic Stimulation (TMS)

Initial motor-cortex mapping was performed using a hand-held MagVenture C-B60 figure-of-eight coil connected to a biphasic MagPro X100 stimulator (MagVenture, Denmark). The optimal motor hotspot for the right first dorsal interosseous (FDI) muscle was identified as the scalp location producing the largest and most consistent visible muscle twitch. Resting motor threshold (rMT) was then determined using the relative-frequency method. The coil was positioned tangentially to the scalp at ≈45° from midline, approximately perpendicular to the central sulcus (Brasil-Neto et al., 1992). Individual stimulation parameters are reported in Table S2.

For the main sessions, all TMS pulses were delivered using a MagVenture Cool-B65 A/P CO figure-of-eight coil connected to a MagPro X100 stimulator. The TMS coil was mounted on and controlled by a robotic positioning arm (TMS Cobot, Axilum Robotics, France), which maintained precise and stable coil positioning throughout the experimental session based on real-time neuronavigation feedback (Richter et al., 2013). This approach minimized position drift and maintained consistent coil-to-cortex distance and orientation across ≈ 1,500 trials. TMS pulses were delivered via an automated interface using the Brain Electrophysiological recording & STimulation (BEST) toolbox (Hassan et al., 2022) and the MAGnetic stimulator Interface Controller (MAGIC) toolbox (Habibollahi Saatlou et al., 2018).

### 2.5. TMS Target Selection and Neuronavigation

The left dlPFC target was defined a priori based on previous work characterizing optimal stimulation sites for reliable early-local TEPs (Parmigiani et al., 2025; Gogulski et al., 2024b). Individual anatomical head models were created using SimNIBS headreco software (Thielscher et al., 2015; Nielsen et al., 2018) applied to each participant’s T1 structural MRI. The standardized dlPFC target (MNI −38, 44, 32) was transformed into native MRI space using nonlinear inverse deformation fields generated with SPM12 (Penny et al., 2011). Coil-angle optimization was performed to minimize early muscle-related artifacts while maximizing EL-TEP amplitude, following real-time optimization procedures (Parmigiani et al., 2025). Real-time neuronavigation used NaviNIBS software (Cline et al., 2025) with 3 OptiTrack PrimeX 41 cameras (NaturalPoint, Corvallis, OR, USA) for six-degree-of-freedom tracking.

### 2.6. Electroencephalography (EEG) Recording

EEG data were continuously recorded using a 64-channel TMS-compatible system with active electrodes (ActiCap, Brain Products GmbH, Germany) and a TMS-compatible amplifier (ActiCHamp Plus, Brain Products GmbH, Germany) sampled at 25 kHz. Electrodes were arranged according to the extended international 10-20 system and referenced online to electrode Cz. Electrode impedances were typically maintained below 5 kΩ throughout the recording session. To minimize off-target auditory and somatosensory artifacts, we implemented a multi-component masking approach. Participants wore insert earbuds delivering continuous broadband white noise which was amplitude-modulated to mask the coil click (Russo et al., 2022), and passive noise-canceling earmuffs (NRR 29 dB) over the earbuds. This combination has been shown to substantially reduce auditory-evoked potentials in TMS-EEG studies (Ross et al., 2022, ter Braack et al., 2015).

### 2.7. TMS-EEG Data Preprocessing

All preprocessing was performed using version 2.0 of the Automated Artifact Removal and TMS-Evoked Potential (AARATEP) pipeline (Cline et al., 2021), which implements a comprehensive sequence of artifact correction procedures optimized for TMS-EEG data. For each experimental block, continuous EEG data were epoched from -800 ms to +1000 ms relative to each TMS pulse. The period from -2 ms to +12 ms surrounding the TMS pulse was replaced with values interpolated using autoregressive extrapolation and blending to remove the large TMS-induced electrical artifact (Cline et al., 2021).

Data were then downsampled to 1 kHz and baseline-corrected using the mean voltage from -500 ms to -10 ms pre-pulse. High-pass filtering above 1 Hz was applied using a modified filtering approach that reduces artifact spread into baseline periods. Bad channels were identified based on quantified noise thresholds and replaced with spatially weighted interpolated values from surrounding electrodes.

Initial per-channel trial averaging was performed using trimmed means, removing the most extreme 10% of voltage values at each time point to reduce the influence of residual artifacts. The first round of independent component analysis (ICA) was performed using FastICA (Hyvärinen, 1999), specifically targeting eye blink artifacts (Mutanen et al., 2020). Components were automatically labeled using the ICLabel classification system (Pion-Tonachini et al., 2019), and components classified as eye blinks with > 90% confidence were removed.

Various non-neuronal noise sources were then attenuated using the source-estimate-utilizing noise-discarding (SOUND) algorithm (Mutanen et al., 2018). Decay artifacts were further reduced using a specialized decay-artifact fitting and removal procedure (Cline et al., 2021). Line noise (60 Hz and harmonics) was attenuated using a 3rd-order Butterworth bandstop filter (58–62 Hz). A second round of ICA was performed with more stringent component rejection criteria targeting any remaining non-neural signals (eye movements, muscle activity, cardiac artifacts) (Rogasch et al., 2014; Pion-Tonachini et al., 2019). The TMS pulse period (-2 to +12 ms) was again interpolated using autoregressive extrapolation and blending (Cline et al., 2021), and data were low-pass filtered below 100 Hz to focus analyses on physiologically relevant frequency ranges. Finally, data were re-referenced to the average of all electrodes.

### 2.8. Quantification of Early-Local TMS-Evoked Potentials (EL-TEPs)

As performed in prior studies, early-local TMS-evoked potentials (EL-TEPs) were extracted from a left frontal region of interest (ROI) consisting of six electrodes (F5, F3, F1, FC5, FC3, FC1) selected to cover the cortical region beneath and immediately surrounding the stimulation site. Previous work has shown that this ROI can capture cortical responses with high reliability (Gogulski et al., 2024a, Gogulski et al., 2024b; Parmigiani et al., 2025).

The EL-TEP was quantified as the peak-to-peak amplitude difference within the 20–80 ms post-stimulation window, with the upper bound extended to 80 ms to capture a positive deflection observed around 55–60 ms post-stimulation (Parmigiani et al., 2025), encompassing the "P30" and "N45" components. These components have neural origins confirmed by intracranial recordings (Keller et al., 2018) and are linked to GABAergic inhibition and glutamatergic excitation in prefrontal microcircuits (Belardinelli et al., 2021; Rogasch et al., 2020; Ahn & Fröhlich, 2021). The peak-to-peak measure captures full response magnitude regardless of small latency differences across participants and conditions.

For each condition, trial-averaged TEPs were computed separately for congruent and incongruent trials using trimmed means. The EL-TEP amplitude for each participant and condition was then calculated by identifying maximum and minimum voltage values within the 20–80 ms window and computing their difference.

### 2.9. Quantification of EL-TEPs from Control Stimulation Sites

To assess the anatomical specificity of cognitive load effects, five participants completed additional blocks with TMS delivered to right dlPFC, left pre-SMA, and vertex control sites. EL-TEPs were defined for each stimulation site as the electrodes nearest to the cortical target: left dlPFC (F5, F3, F1, FC5, FC3, FC1), right dlPFC (F6, F4, F2, FC6, FC4, FC2), left pre-SMA (C1, C3, C5, CP1, CP3, CP5), and vertex (CPz, Pz, POz, P1, P2). (Gogulski et al., 2024b; Rosanova et al., 2009, Casarotto et al., 2010 Parmigiani et al., 2025). Each control site underwent identical EL-TEP quantification (20–80 ms window) following the real-time optimization procedures (Parmigiani et al., 2025). ‘Downstream’ dlPFC responses following stimulation of each control site were also extracted to evaluate whether cognitive load effects were specific to the left dlPFC stimulation site. Here, following TMS to each non-dlPFC site, we extracted the TEP in the left prefrontal region (F5, F3, F1, FC5, FC3, FC1). These control sites were included to determine whether cognitive load effects on local EL-TEPs were specific to the left dlPFC, or whether similar modulation occurred at other stimulation sites.

### 2.10. Quantification of Midfrontal Theta Rhythm Power

Theta oscillations (4–8 Hz) were quantified from a midfrontal site (FCz) after applying a surface Laplacian (Hjorth nearest-neighbor transform) using Fz, Cz, FC1, and FC2 as the reference ring, consistent with established conflict-related theta sources (Cavanagh & Frank, 2014; Cohen, 2014). EEG epochs spanning -250 to +250 ms relative to MSIT stimulus onset captured stimulus encoding and conflict processing prior to TMS. Power spectra were computed using FFT with a Hanning window (0.5 Hz resolution) (Welch, 1967). To isolate oscillatory activity from the aperiodic 1/f background, spectra were log-transformed, a linear fit was estimated outside the theta band, and this was subtracted from the original spectrum following established methods (Donoghue et al., 2020). Theta power was quantified as the mean 4–8 Hz power from the 1/f-corrected spectrum.

Theta phase at the time of each TMS pulse was estimated offline from the last 500 ms of the pre-stimulus EEG signal (Figure 4A), band-pass filtered with a two-pass (zero-phase) FIR filter (4–8 Hz), and converted to instantaneous phase via the Hilbert transform. Because the phase at which TMS was applied could not be directly calculated due to signal corruption by TMS-related artifacts and evoked potentials, phase was estimated for the uncorrupted time point exactly one individual theta cycle earlier (Hassan et al., 2025; Bergmann et al., 2019).

### 2.11. Behavioral Data Analysis

Behavioral performance was assessed using accuracy and reaction time. Trial-level data were aggregated to block-level means per participant and averaged within cognitive interference conditions. Trial-type–specific analyses were also conducted to separate congruency and block-level effects. Reaction time variability was quantified as within-block standard deviation. These approaches align with cognitive-control analyses in prior MSIT and conflict paradigms (Bush et al., 2003; Bush & Shin, 2006).

### 2.12. Statistical Analyses

Statistical analyses were performed using linear mixed-effects (LME) models implemented in Python’s statsmodels package (Bates et al., 2015). LME models were chosen for their ability to handle nested structures, unbalanced data, and subject-level variability. Separate LME models were constructed for reaction time, accuracy, EL-TEP amplitude, and theta power as dependent variables.

#### Behavioral Analyses

Separate LME models were constructed for reaction time and accuracy as dependent variables. For both models, cognitive interference condition (0-back control, 0%, 20%, 33%, 66%, 80%, 100% incongruent) was included as a fixed effect, with participants included as random intercepts. Post-hoc pairwise comparisons were conducted using estimated marginal means with Bonferroni correction for multiple comparisons (Lenth, 2019).

Based on pilot data suggesting peak behavioral disruption at intermediate interference levels rather than at maximum incongruency, we hypothesized that cognitive load would peak at intermediate interference levels where trial-type uncertainty is greatest, rather than at maximum incongruency where participants can adopt fixed response strategies. To test this, we conducted focused contrasts comparing the 33% congruent condition, which maximized trial-type uncertainty, to conditions with both lower interference (0-back, 100%, 80% congruent) and higher incongruency proportions (20%, 0% congruent) where fixed response strategies could be adopted. This contrast tests whether cognitive load, as defined by behavioral performance, peaks at intermediate interference levels.

#### EL-TEP Modulation Analyses

The primary analysis examined whether EL-TEP amplitudes varied systematically with cognitive load. An LME model was constructed with EL-TEP amplitude as the dependent variable and the following fixed effects: cognitive interference condition (7 levels), trial type (congruent vs. incongruent), and their interaction. Participant was included as a random intercept. Post-hoc tests examined specific contrasts of interest, including: (1) differences between each condition (7 levels: control [0-back], and 6 MSIT conditions with 100%, 80%, 66%, 33%, 20%, and 0% congruent trials), (2) differences between congruent and incongruent trials within each of the four mixed-load MSIT conditions (80%, 66%, 33%, and 20% congruent), and (3) the shape of the cognitive interference-response curve.

#### Spatial Specificity Analyses

To assess whether cognitive interference effects were specific to the left dlPFC stimulation site, we constructed a hierarchical LME model including data from all four stimulation sites (left dlPFC, right dlPFC, pre-SMA, vertex). This model included fixed effects of cognitive interference, stimulation site, trial type, and two-way interactions between MSIT conditions and stimulation site, and cognitive MSIT conditions and trial type. Participant was included as a random intercept. Separate follow-up LME models were conducted for each stimulation site independently to characterize site-specific effects.

#### Multiple Comparisons Correction

All post-hoc pairwise comparisons were corrected for multiple comparisons using Bonferroni correction. The criterion for statistical significance was set at α = 0.05 for all tests (two-tailed).

#### Effect Sizes

For pairwise comparisons, effect sizes were quantified using Cohen’s d, calculated using the mean difference divided by the pooled standard deviation. Effect size interpretations followed standard conventions: small (d = 0.2), medium (d = 0.5), and large (d = 0.8) (Cohen, 1988).

Throughout the manuscript, group data are presented as mean ± standard error of the mean (SEM) unless otherwise specified. Individual participant data are shown in supplementary figures to illustrate effect consistency and individual variability. All statistical values are reported with specific p-values when p > 0.001; for smaller p-values, we report p < 0.001.

## 3. Results

### 3.1. Reaction times and accuracy scale with cognitive interference level

#### 3.1.1. Reaction times and accuracy show systematic modulation with cognitive interference

In the main experiment (N=27), we first investigated how cognitive interference, manipulated by varying the ratio of congruent to incongruent trials, affected reaction times and accuracy on the multi-source interference task (MSIT). Participants performed the MSIT under seven conditions with concurrent TMS: a control condition equivalent to a 0-back task, and six MSIT conditions with 100%, 80%, 66%, 33%, 20%, and 0% congruent trials corresponding to increasing proportions of incongruent trials. Participants also completed no-TMS blocks and additional control blocks (see Methods), but the present analyses focus on the seven TMS-concurrent conditions. Analysis using linear mixed-effects (LME) models revealed a significant main effect of cognitive interference condition on reaction times (Bonferroni-corrected pairwise comparisons: all pairs p < 0.05, with 19 of 21 comparisons p < 0.001, Cohen’s d range: 1.05–9.23; see significance matrix in Figure 2C. Similarly, we observed a significant main effect of cognitive interference on task accuracy (16 of 21 pairwise comparisons p < 0.001, 2 p < 0.05, 3 n.s., Cohen’s d range: 0.86–6.27; see Figure 2D), indicating that accuracy systematically varied with cognitive interference level (Figure 2B, 2D).

**Figure 2:**
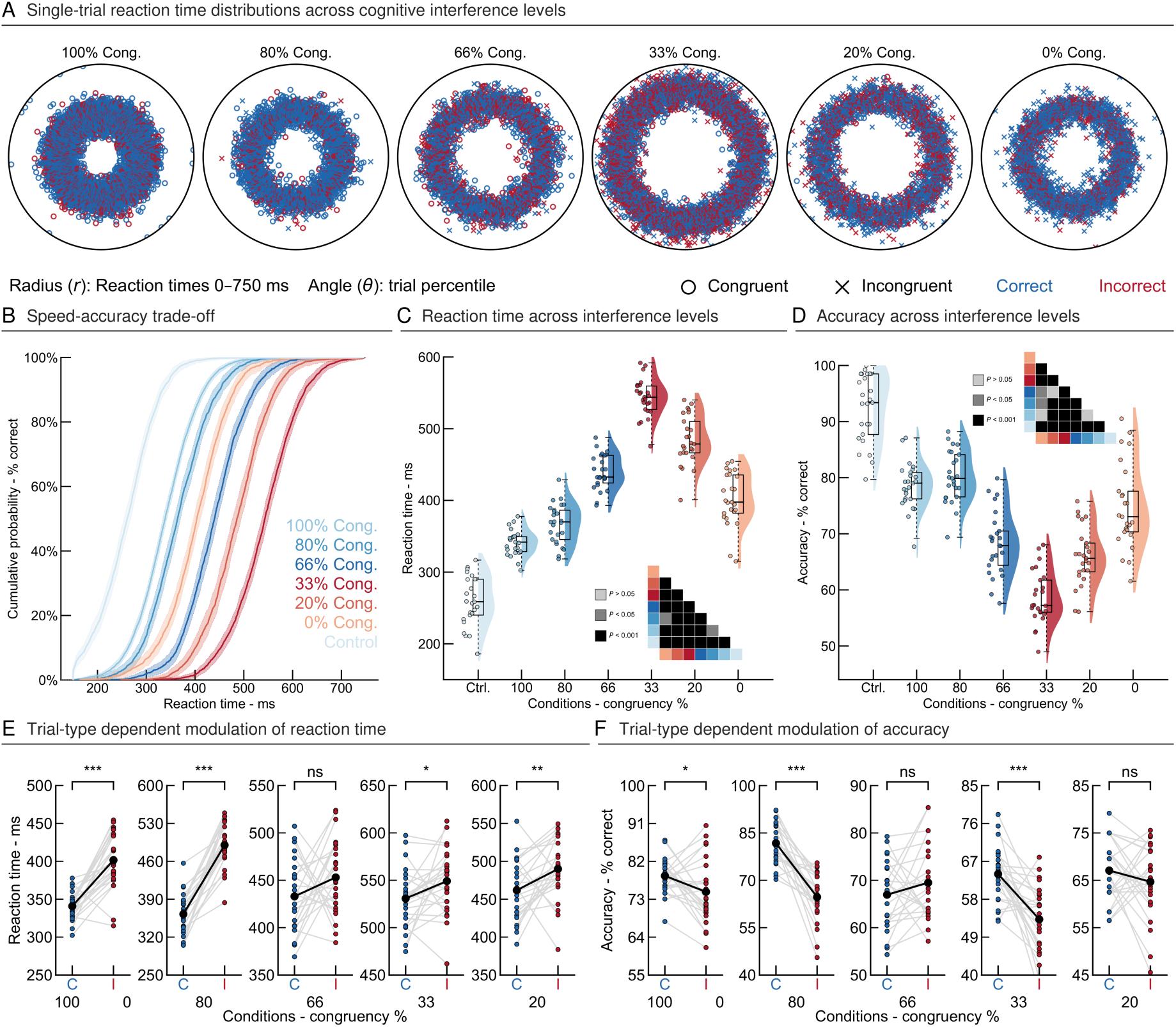
Behavioral validation of cognitive interference manipulation during the Multi-Source Interference Task. (A) Single-trial reaction time distributions across cognitive interference conditions (N=27). Circular plots display individual trial reaction times (radius: 0-750 ms) as a function of trial position (angle: trial percentile). Blue color represent correct, red represent inorrect trials. Circle represent congruent, X represent incongruent trials. (B) Speed-accuracy tradeoff curves showing cumulative probability of correct responses as a function of reaction time across all cognitive interference conditions. Curves shift rightward with increasing load (blue to red gradient), indicating slower responses required to maintain accuracy under higher cognitive load. (C) Group-level reaction times across conditions. Violin plots show reaction time distributions for each condition. Dots represent individual participants. Significance matrix (bottom right) indicates pairwise comparisons. (D) Group-level accuracy across conditions following same format as panel C. (E) Trial-type dependent modulation of reaction times. Five paired comparisons show reaction times for congruent (C, blue) versus incongruent (I, red) trials within each mixed-load condition and a fully congruent (100%) vs fully incongruent (0%) condition. Gray lines connect data from individual participants. Incongruent trials consistently showed longer average reaction times than congruent trials across all conditions (***P < 0.001, **P < 0.01, *P < 0.05). (F) Trial-type dependent modulation of accuracy. Six paired comparisons show accuracy for congruent versus incongruent trials, following same format as panel E. Congruent trials showed consistently higher average accuracy than incongruent trials (***P < 0.001, **P < 0.01, *P < 0.05). Together, these behavioral results confirm that the MSIT successfully manipulated cognitive load, with robust interference effects evident in both reaction time and accuracy measures.

#### 3.1.2. Reaction times exhibit non-linear relationship with cognitive interference

Analysis of reaction time data revealed a non-linear relationship with cognitive interference (Figure 2C). Reaction times significantly increased from the control (0-back) condition (260±7 ms) through intermediate conditions (100% congruent: 341±4 ms; 80% congruent: 367±6 ms; 66% congruent: 440±5 ms) to peak at the 33% congruent condition (543±5 ms), representing the highest response latency. As the proportion of incongruent trials further increased, reaction times decreased compared to the 33% congruent condition (20% congruent: 484±7 ms; 0% congruent: 402±7 ms). Pairwise LME comparisons with Bonferroni correction confirmed significant differences between adjacent conditions (all p < 0.01). Within-participant reaction time variability (computed as within-block standard deviation) also showed significant modulation across conditions, with maximum variability observed at intermediate incongruency levels (not shown). Within each condition, incongruent trials showed longer reaction times, reaching significance at 80% (p < 0.001), 33% (p = 0.028), and 20% congruent (p = 0.009), though not at 66% congruent (p = 0.065) (Figure 2E). The between-block comparison of the purely congruent (100%) and purely incongruent (0%) conditions was also significant (p < 0.001) (Figure 2E), confirming the robust interference effect of the MSIT paradigm.

#### 3.1.3. Task accuracy follows a similar non-linear pattern with cognitive interference

Paralleling the reaction time results, task accuracy also exhibited a non-linear relationship with cognitive interference (Figure 2D). Accuracy was highest in the control condition (92.4±1.2%) and decreased with increasing incongruency proportion through intermediate conditions (100% congruent: 78.6±0.8%; 80% congruent: 80.2±0.9%; 66% congruent: 68.2±1.1%), reaching a minimum at the 33% congruent condition (58.6±0.9%). At higher incongruency proportions, accuracy partially recovered (20% congruent: 65.9±1.0%; 0% congruent: 74.8±1.5%), mirroring the non-linear pattern observed in reaction times. Within each condition, incongruent trials showed lower accuracy, reaching significance at 100% (p = 0.021), 80% (p < 0.001), and 33% congruent (p < 0.001), though not at 66% (p = 0.191) or 20% congruent (p = 0.233) (Figure 2F). The between-block comparison of the purely congruent (100%) and purely incongruent (0%) conditions was also significant (p = 0.021) (Figure 2F), further confirming the effectiveness of manipulating cognitive effort (see Figure S1 for individual subject distributions).

### 3.2. Left prefrontal excitability tracks cognitive load

#### 3.2.1. Prefrontal EL-TEPs are modulated with varying cognitive load levels

EL-TEP amplitudes from left prefrontal cortex scaled systematically with cognitive load level. Here, cognitive load is defined by behavioral performance, with peak load occurring at 33% congruency, where trial-type uncertainty was highest and accuracy and reaction times were worst (Figure 2A-D). Grand average butterfly plots with corresponding topographical voltage distributions at relevant latencies (40ms, 55ms, and 70ms) illustrate the spatiotemporal dynamics of early TEPs (Figure 3A). These topographical maps showed prominent voltage deflections over left frontal electrodes, consistent with local cortical activation following left dlPFC stimulation. We next quantified early local TEPs (EL-TEPs) to capture this early local activation (see Methods). Grand averaged EL-TEPs at control and parametrically varying cognitive load revealed a significant main effect of cognitive load on prefrontal EL-TEP amplitudes (N=27, pairwise LME with Bonferroni correction, Cohen’s d = 1.55–4.12 vs. control; Figure 3B). Pairwise comparisons demonstrated that EL-TEP amplitudes significantly increased from the control condition (1.58±0.16 μV) through the 100% (2.34±0.16 μV, p <0.01), 80% (2.94±0.17 μV, p<0.001), 66% (3.40±0.17 μV, p<0.001), 33% (4.09±0.19 μV, p<0.001), and 20% congruent conditions, with the 20% congruent condition showing the largest amplitude (4.61±0.19 μV, p<0.001). The 0% congruent condition (3.34±0.16 μV) showed a decrease from the 20% condition, and this difference was significant (p < 0.001). The cognitive load vs. EL-TEP amplitude curve revealed a significant non-linear relationship between increasing cognitive interference and neurophysiological response (Figure 3B), similar to the non-linear behavioral findings on reaction time and accuracy (Figure 2C, 2D). Grand-average EL-TEP waveforms stratified by reaction time windows further demonstrated that trials with longer reaction times were associated with progressively larger EL-TEP amplitudes (Figure 3C), and condition-wise individual subject means revealed a positive relationship between reaction time and EL-TEP amplitude (Figure 3D). See Figure S3 for comparison of EL-TEP vs spontaneous EEG amplitudes, Figure S4 for individual subject traces, and Figure S5 for butterfly plots.

**Figure 3:**
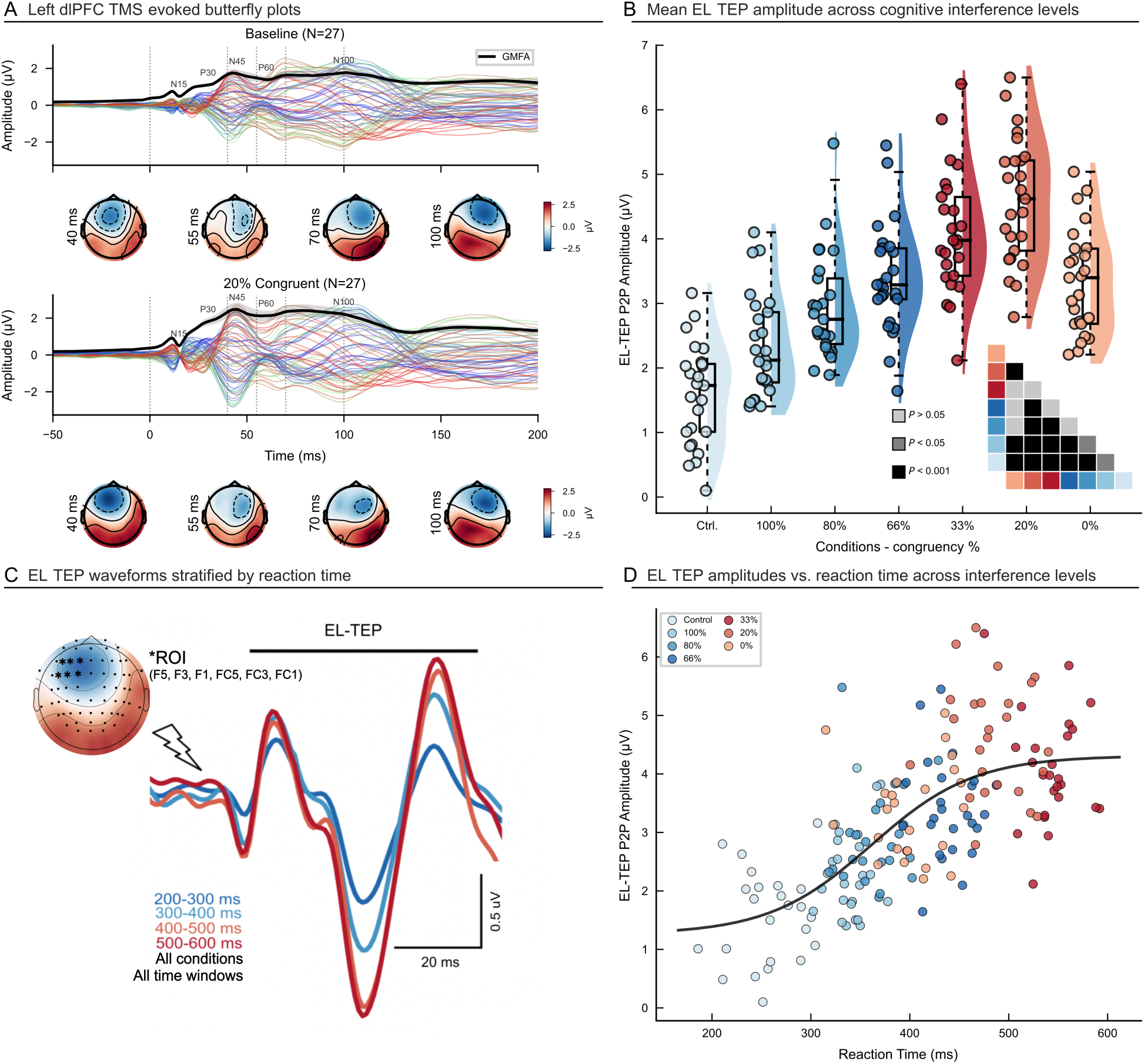
Cognitive interference modulates prefrontal early local TMS-evoked potentials (EL-TEPs) as a function of task congruency and behavioral performance. A) Grand-average butterfly plots of TMS-evoked EEG responses (TMS delivered at t = 0 ms) recorded over the left dorsolateral prefrontal cortex (dlPFC) under baseline and MSIT 20% congruency conditions, illustrating distinct spatiotemporal response profiles. Insets show corresponding scalp topographies at four representative latencies (40, 55, 70 and 100 ms) for each condition, highlighting focal activation differences across the cortical surface. B) Violin–box plots showing EL-TEP amplitudes across control and parametrically varying cognitive interference levels (100– 0% congruency). Increasing incongruency produced systematic amplitude increases, with the inset significance matrix indicating post hoc contrasts (p < 0.05; p < 0.001). C) Grand-average EL-TEP waveforms (N = 27) grouped by reaction time windows (200–300 ms, 300–400 ms, 400–500 ms, 500–600 ms), demonstrating that trials with longer reaction times are associated with progressively larger EL-TEP amplitudes. D) Scatter plot of condition-wise individual subject means illustrating a nonlinear positive relationship between reaction time and EL-TEP amplitude, with color coding reflecting cognitive interference level (congruency level). Together, these findings show that prefrontal cortical excitability, indexed by EL-TEPs, scales with both cognitive interference level and behavioral slowing, indicating a state-dependent modulation of TMS-evoked responses by cognitive control demands.

### 3.3. Midfrontal theta power parallels prefrontal excitability across conditions

#### 3.3.1. Theta power scales with cognitive load

Midfrontal theta power showed a load-dependent response pattern that paralleled the prefrontal excitability profile. Of note, theta power was quantified from the pre-TMS time window (−500 to 0 ms before the TMS pulse, corresponding to −250 to +250 ms relative to MSIT stimulus onset; Figure 4A) to ensure that TMS-evoked neural activity did not contaminate theta power estimates. After applying 1/f correction to the power spectrum (see Methods; Figure 4A) of the peri-MSIT visual stimulus period (-250 to 250ms relative to stimulus onset), we observed a non-linear modulation of theta power across MSIT conditions (Figure 4B, 4H). Linear mixed-effects modeling (N=27) revealed a significant main effect of cognitive interference on theta power (pairwise LME with Bonferroni correction: 13 of 15 comparisons p < 0.05; see significance matrix in Figure 4H). Scalp topographies and time-frequency representations confirmed this effect was localized to midfrontal sites (Figure 4E). Theta power increased from the 100% congruent condition (0.211±0.004 µV²/Hz) through the 80% (0.293±0.006 µV²/Hz) condition, with the highest theta power at 33% congruent (0.406±0.004 µV²/Hz; all p<0.001 after Bonferroni correction). The 66% condition showed intermediate theta power (0.321±0.005 µV²/Hz). At higher incongruency proportions, theta power decreased from the 33% peak (20% congruent: 0.320±0.005 µV²/Hz; 0% congruent: 0.294±0.006 µV²/Hz) (Figure 4H), mirroring the non-linear pattern observed in reaction times (Figure 2C) and EL-TEP amplitudes (Figure 3B). Comparison of no cognitive load versus cognitive load conditions revealed significantly higher theta power during load (no-load: 0.097±0.006 µV²/Hz; load: 0.305±0.013 µV²/Hz; LME p < 0.001, Cohen’s d = 4.02; Figure 4F). Theta power did not differ across control conditions (baseline, control task, and pre-stimulus time windows; all p > 0.05; Figure 4G), confirming that the observed modulation was specific to active cognitive engagement. (see Figure S2 for timing control analyses)

**Figure 4.**
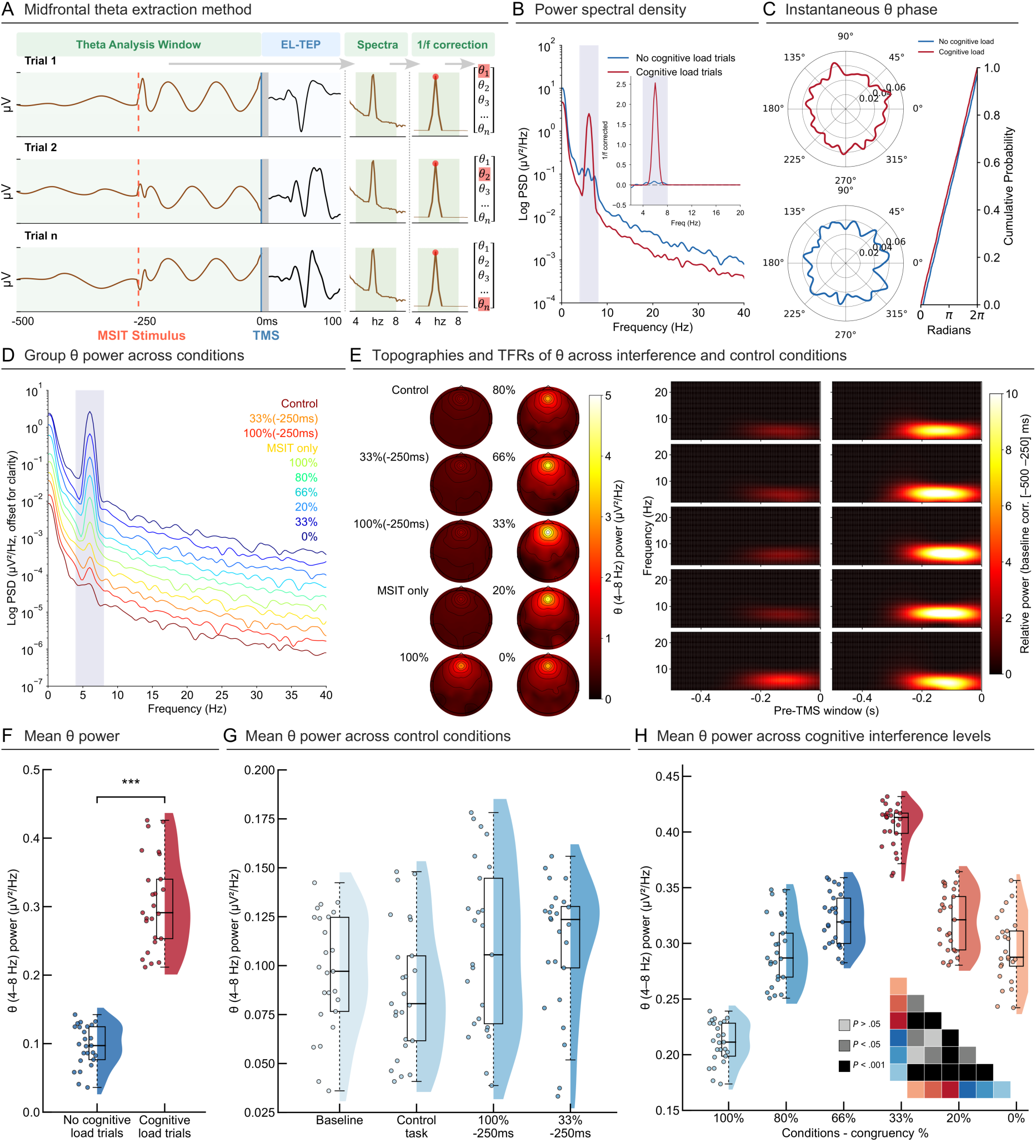
Midfrontal theta power increases with cognitive load. (A) Schematic of the analysis: theta (4–8 Hz) is extracted from pre-stimulus EEG (−500 to 0 ms before TMS pulse). and the 1/f background is removed. This window corresponds to −250 to +250 ms relative to MSIT visual stimulus onset for all MSIT task conditions (TMS delivered at +250 ms post-stimulus), and to −750 to −250 ms relative to MSIT visual stimulus onset for the two control conditions (TMS delivered at −250 ms pre-stimulus) (B) Power spectra (FOOOF-corrected) show higher theta amplitude during cognitive-load trials. (C) Circular phase plots and cumulative phase distributions reveal no load-dependent instantaneous phase differences. (D) Group power spectra across load levels display a monotonic increase in theta with greater interference. (E) Scalp maps and time–frequency plots localize this effect to midfrontal sites and show stronger theta under higher load. TFRs are baseline corrected from [-500 to -250 ms] using the relative power method. (F) Violin plot: mean theta power is greater in cognitive-load vs no-load trials (***p < 0.001). (G) Theta power across baseline, control, and early stimulation MSIT conditions. (H) Graded increase in theta power across all cognitive interference series (from 100% to 0% congruent). Significance matrix (bottom right) indicates pairwise comparisons.

### 3.4. Load-dependent excitability is specific to left prefrontal cortex

#### 3.4.1. Behavioral performance remains consistent across stimulation sites

To determine whether the sensitivity of EL-TEPs to cognitive load was specific to the left prefrontal cortex or a more general phenomenon, in a separate control experiment, we applied TMS pulses to four cortical regions in an independent cohort of participants (N=5) and measured EL-TEPs locally in those cortical regions. Reaction times at three cognitive interference conditions (MSIT Control Task, 100% congruent trials, 33% congruent trials) were measured across four TMS sites: left dlPFC (LdlPFC), right dlPFC (RdlPFC), left pre-SMA (LSMA) and vertex (Figure 5A-B). Pairwise LME comparisons with Bonferroni correction revealed significant differences in reaction times and accuracy across all cognitive interference conditions at each stimulation site (all pairwise p < 0.001; Figure 5A-B), with similar behavioral patterns across sites indicating that the cognitive interference manipulation impacted behavior consistently regardless of stimulation site.

**Figure 5:**
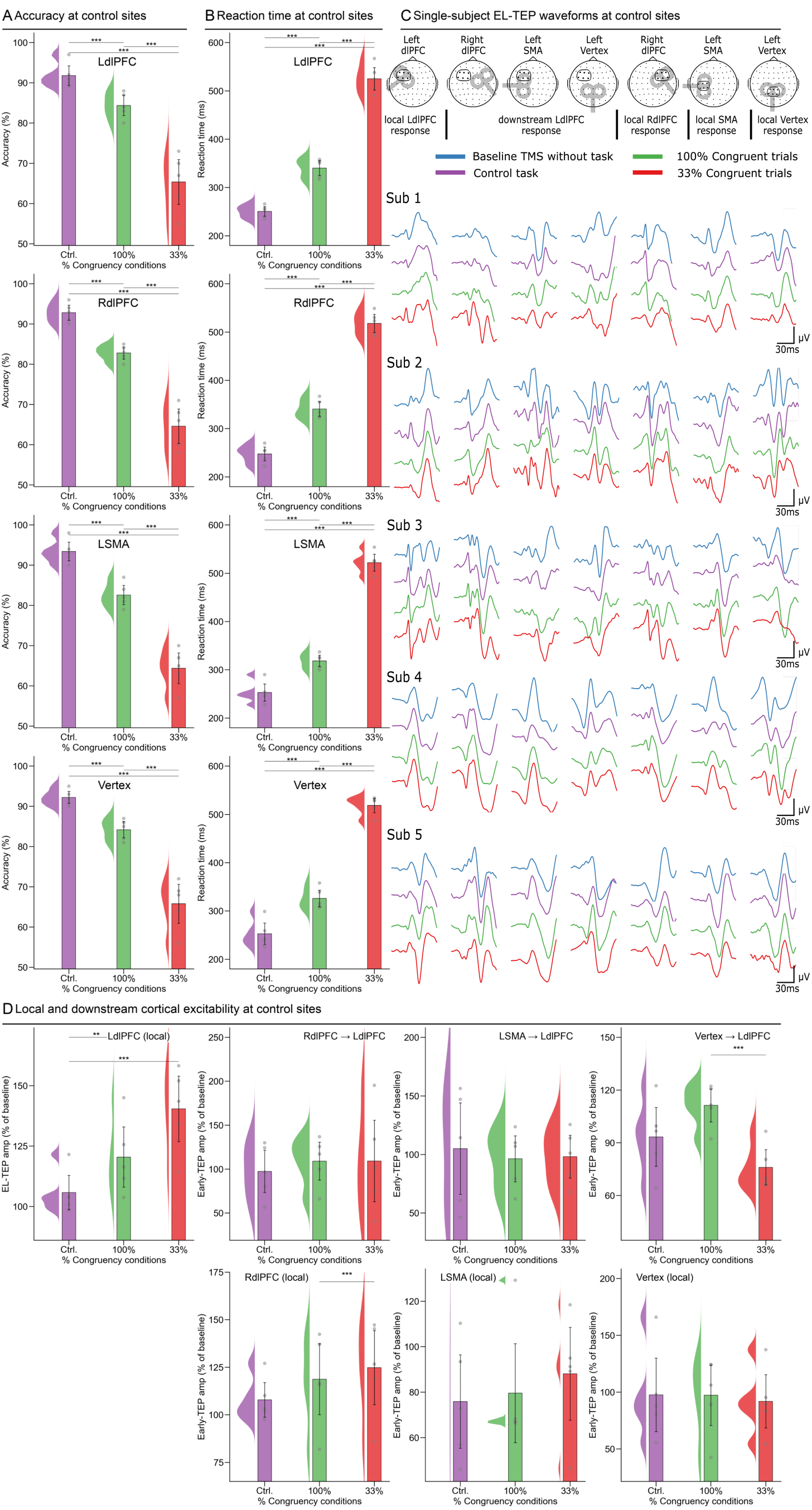
Cognitive interference does not modulate EL-TEPs in right dlPFC, pre-SMA and vertex. A) Reaction times at three cognitive interference conditions (MSIT Control Task, 100% MSIT Congruent trials, 33% MSIT Congruent trials) across four stimulation sites: left dlPFC (LdlPFC), right dlPFC (RdlPFC), left pre-SMA (LSMA) and vertex (N=5). B) Mean reaction time comparison between the three cognitive interference conditions (MSIT Control Task, 100% MSIT Congruent trials, 33% MSIT Congruent trials) for each stimulation site: left dlPFC (LdlPFC), right dlPFC (RdlPFC), left pre-SMA (LSMA) and vertex (N=5). C) Individual EL-TEP/TEP (20-80ms) traces recorded locally for LdlPFC, RdlPFC, LSMA, and Vertex as well as downstream LdlPFC responses following RdlPFC, LSMA and Vertex stimulation. D) EL-TEP amplitude quantified as percentage from baseline TEP condition measured locally for LdlPFC, RdlPFC, LSMA, and Vertex as well as downstream LdlPFC responses following RdlPFC, LSMA and Vertex stimulation. Significance of post hoc comparisons is indicated as follows: ∗p < 0.05; ∗∗p < 0.01; ∗∗∗p < 0.001.

#### 3.4.2. Site-specific differences in EL-TEP modulation across cognitive interference conditions

We observed distinct pattern differences within individual EL-TEP traces recorded locally after TMS pulses were applied to LdlPFC, RdlPFC, LSMA, and vertex, as well as downstream LdlPFC responses following RdlPFC, LSMA and vertex stimulation (Figure 5C). While left dlPFC TMS produced EL-TEPs that varied systematically with cognitive load, similar modulations were not observed for the other stimulation sites (Figure 5D). Pairwise LME comparisons with Bonferroni correction showed significant load-dependent EL-TEP modulation for left dlPFC TMS (Control: 105.79±3.63% baseline; 100% congruent: 120.47±6.34%; 33% congruent: 140.42±6.90%; Ctrl vs 100%: p = 0.007; Ctrl vs 33%: p < 0.001). In contrast, EL-TEPs elicited from right dlPFC (local: Control: 107.88±4.66%; 100% congruent: 118.74±9.52%; 33% congruent: 124.80±9.92%), pre-SMA (local: Control: 75.90±10.50%; 100% congruent: 79.60±11.11%; 33% congruent: 88.07±10.42%), and vertex (local: Control: 97.61±16.53%; 100% congruent: 97.30±13.56%; 33% congruent: 91.97±11.95%) showed no consistent load-dependent modulation (pre-SMA and vertex: all pairwise p > 0.05; right dlPFC 100% vs 33%: p < 0.001, other comparisons p > 0.05). Similarly, downstream LdlPFC TEPs following RdlPFC, LSMA, and Vertex TMS also showed no consistent modulation by cognitive load. Downstream LdlPFC responses: RdlPFC→LdlPFC (Control: 97.50±12.28%; 100% congruent: 109.15±11.05%; 33% congruent: 109.29±23.65%; all p > 0.05), LSMA→LdlPFC (Control: 104.98±19.89%; 100% congruent: 96.37±10.02%; 33% congruent: 98.18±9.34%; all p > 0.05), and Vertex→LdlPFC (Control: 93.36±8.55%; 100% congruent: 111.29±4.79%; 33% congruent: 76.05±5.12%; 100% vs 33%: p < 0.001, other comparisons p > 0.05).

## 4. Discussion

### Prefrontal excitability dynamically varies with cognitive load

Here we demonstrate that prefrontal cortical excitability, measured through TMS-evoked potentials, provides a window into cognitive state, potentially offering a long-sought bridge between basic neuroscience and clinical application. Prefrontal EL-TEP amplitudes systematically scaled with cognitive interference, peaking at moderate interference levels before decreasing at maximal interference levels (Figure 3B). This non-linear curve aligned with behavioral performance (Figure 2C-D, Figure 3D), suggesting these EL-TEPs scale with cognitive load. This cognitive state-dependent modulation was anatomically specific to the left prefrontal cortex, as EL-TEPs from the right prefrontal cortex, presupplementary motor area, and vertex showed no reliable interference effects (Figure 5). Changes in prefrontal excitability were paralleled by modulations in midfrontal theta power (Figure 3B, Figure 4H), a well-established marker of cognitive control. These findings suggest intrinsic prefrontal theta and prefrontal excitability measures may represent the same underlying prefrontal network state, complementing prior theta oscillation and fMRI findings (Cavanagh & Frank, 2014; Bush & Shin, 2006). Together, these findings reveal that prefrontal excitability fluctuates in tight coordination with cognitive load and theta power, suggesting that EL-TEPs can serve as a temporally precise marker of dynamic brain state changes during cognitive control.

### Prefrontal excitability measures as cognitive load markers

Here we demonstrate that prefrontal EL-TEPs systematically scale with cognitive load during the MSIT, providing a direct neurophysiological readout of cognitive control engagement. EL-TEP amplitudes increased parametrically with cognitive interference levels, peaking at intermediate incongruency levels (20–33% congruent) before showing a slight decrease under maximal interference (Figure 3B). This pattern may reflect instantaneous gain control in prefrontal circuits – a mechanistic signal of how local inhibitory-excitatory balance adjusts to meet cognitive load (Murray et al., 2014; Premoli et al., 2014). Our findings extend previous work establishing EL-TEPs as robust and reliable measures of prefrontal excitability across different stimulation sites (Gogulski et al., 2024a; Gogulski et al., 2024b), and optimizable through real-time parameter adjustment (Parmigiani et al., 2025). The load-dependent EL-TEP modulation observed here was anatomically specific to prefrontal regions, as both TEPs recorded at the stimulation site and at prefrontal electrodes after TMS to non-prefrontal sites showed no reliable load effects. Notably, EL-TEPs at the pre-stimulus TMS time point (−250 ms) showed no cognitive load modulation, confirming that the effect is time-locked to active cognitive processing (Figure S2). The rapid measurement capability of EL-TEPs opens new possibilities for translational applications: real-time tracking of prefrontal excitability could enable closed-loop control systems that optimize stimulation timing based on instantaneous brain states, potentially enhancing therapeutic efficacy by delivering TMS when prefrontal circuits are most responsive. The ability to track dynamic changes in prefrontal engagement during cognitive control adds important validation to EL-TEPs as physiological readouts of circuit-level dynamics.

### Neural mechanisms underlying state-dependent prefrontal excitability

The ability of EL-TEPs to track cognitive load provides insights into the circuit mechanisms underlying prefrontal cognitive control. The early latency (20-80ms) of these responses suggests short-latency engagement of local inhibitory and excitatory circuits, potentially reflecting a combination of GABA-mediated inhibition and NMDA receptor-dependent excitation in prefrontal microcircuits (Belardinelli et al., 2021). The initial positive deflection (20-30ms) likely reflects fast GABAergic inhibition from parvalbumin-positive interneurons, which are known to shape cortical excitability and gamma-theta coupling during cognitive tasks (Sohal et al., 2009; Cardin et al., 2009). The subsequent negative component (30-50ms) may arise from NMDA receptor-dependent recurrent excitation within layer 2/3 pyramidal networks, which show prolonged depolarization following stimulation (Rogasch et al., 2020; Belardinelli et al., 2021). The final positive peak (60-80ms) potentially reflects feedback inhibition from somatostatin-positive interneurons, which regulate gain control in prefrontal circuits during cognitive processing (Abbas et al., 2018; Urban-Ciecko & Bhatt, 2016). These early evoked potentials generally align temporally with intracranial recordings showing that direct electrical stimulation evokes early responses within 60ms (Keller et al., 2018; Wang et al., 2024). We observed EL-TEPs that showed a non-linear state dependency pattern, peaking at intermediate incongruency levels before declining at maximal incongruency. This pattern is consistent with reduced cognitive control demands at extreme block compositions where trial-type uncertainty is minimal and participants can adopt fixed response strategies. The anatomical specificity of these effects to dorsolateral prefrontal cortex, with minimal modulation at non-prefrontal sites, supports engagement of the well-characterized cognitive control network centered on dorsolateral prefrontal and anterior cingulate cortices (Bush & Shin, 2006). Moreover, the parallel modulation of EL-TEP amplitudes and theta power suggests that TMS may be probing the same prefrontal circuits that naturally synchronize in the theta band during cognitive control (Smith et al., 2019). These findings advance understanding of the neurophysiological basis of cognitive control, indicating that prefrontal excitability is dynamically regulated based on cognitive load through rapid modulation of local circuit properties.

### Theta oscillations as a bridge to cognitive control networks

Our findings reveal convergent modulation of prefrontal excitability and theta oscillations during cognitive control. EL-TEP amplitudes and theta power showed parallel modulation patterns across cognitive interference conditions (Figure 3B, Figure 4H), with both measures showing increases during higher cognitive load states. This relationship was specific to the prefrontal cortex and scaled systematically with task demands, suggesting these measures reflect converging indices of the same cognitive control network state. Previous work has shown that theta oscillations coordinate prefrontal networks during conflict monitoring (Cavanagh & Frank, 2014), response inhibition (Zavala et al., 2018), and error processing (Cavanagh & Shackman, 2015), potentially reflecting or enabling underlying changes in prefrontal circuit excitability. By linking causal excitability probes (EL-TEPs) to intrinsic theta oscillations, our findings provide evidence that cognitive control emerges through coordinated changes in both prefrontal excitability and network synchronization, supporting a mechanistic model in which theta oscillations and prefrontal excitability are co-modulated during cognitive control. Given that theta phase organizes neural processing into alternating windows of enhanced and reduced excitability, this framework generates testable predictions about the phase-dependent modulation of cortical excitability and offers new avenues for investigating how theta-mediated excitability dynamics contribute to flexible cognitive control (Cavanagh & Frank, 2014; Cohen & Donner, 2013; Smith et al., 2019; Helfrich & Knight, 2016).

### Limitations and future directions

Several limitations must be considered. A key conceptual limitation is that while we capture excitability snapshots at discrete moments, our approach does not measure directional connectivity or effective connectivity between prefrontal regions and broader cognitive control networks. This leaves open questions about how state-dependent excitability changes propagate through distributed circuits and whether excitability modulations in dlPFC causally influence downstream network dynamics or reflect upstream inputs from other nodes in the control network. While our sample size (N=27) is modest, the robust single-subject effects (Figure S1, S4) and anatomical specificity demonstrated through control site stimulation establish the reliability of the phenomenon. Technical challenges inherent to TMS-EEG, including limited spatial resolution and potential sensory effects, were mitigated through our multi-component masking approach and control conditions, though individual variability in TMS responses remains an important consideration for future work. The fixed +250ms latency between stimulus onset and TMS pulse may not capture the full temporal dynamics of cognitive state changes and warrants further exploration. Additionally, our focus on a single cognitive paradigm (MSIT) leaves open questions about whether these effects generalize to other forms of cognitive control. Finally, while we demonstrate these effects in healthy participants, validation in clinical populations is needed to establish therapeutic relevance, particularly given potential differences in prefrontal circuit dynamics in psychiatric conditions.

This work also suggests potential clinical applications, though these require systematic validation. Future clinical trials could assess whether performing cognitive control tasks during therapeutic TMS sessions enhances efficacy through state-dependent plasticity mechanisms. Longitudinal studies examining how cognitive-state dependent EL-TEPs change over the course of treatment could yield valuable biomarkers for monitoring therapeutic response. Testing whether baseline excitability dynamics predict treatment outcomes and whether real-time excitability tracking could guide adaptive stimulation represents a promising direction for optimizing brain stimulation therapies. If validated and translated to patient populations, state-dependent excitability tracking could transform TMS from a static intervention into an adaptive, brain-state-responsive therapy. Together, these results establish prefrontal excitability as a real-time, regionally selective physiological readout of cognitive state, one that may ultimately guide when and where to stimulate.

## Acknowledgements.

We extend gratitude to all of our research participants. We would also like to acknowledge the generous contributions of the members of the Personalized Neurotherapeutics Laboratory for helpful feedback on the manuscript and throughout the course of the study. This research was supported by the National Institute of Mental Health under award number R01MH126639, R01MH139650, R01MH129018, and a Burroughs Wellcome Fund Career Award for Medical Scientists (CJK). UH was supported by the Sleep Research Society Foundation, Stanford Spark and Dean’s fellowship programmes. JMR was supported by the Department of Veterans Affairs Office of Academic Affiliations Advanced Fellowship Program in Mental Illness Research and Treatment, the Medical Research Service of the Veterans Affairs Palo Alto Health Care System and the Department of Veterans Affairs Sierra-Pacific Data Science Fellowship. JG was supported by personal grants from Orion Research Foundation, the Finnish Medical Foundation, and Emil Aaltonen Foundation.

## Declaration of Interest

CJK holds equity in Alto Neuroscience, Inc. and Flow Neuroscience, Inc.

## Supplementary Materials

**Figure S1:**
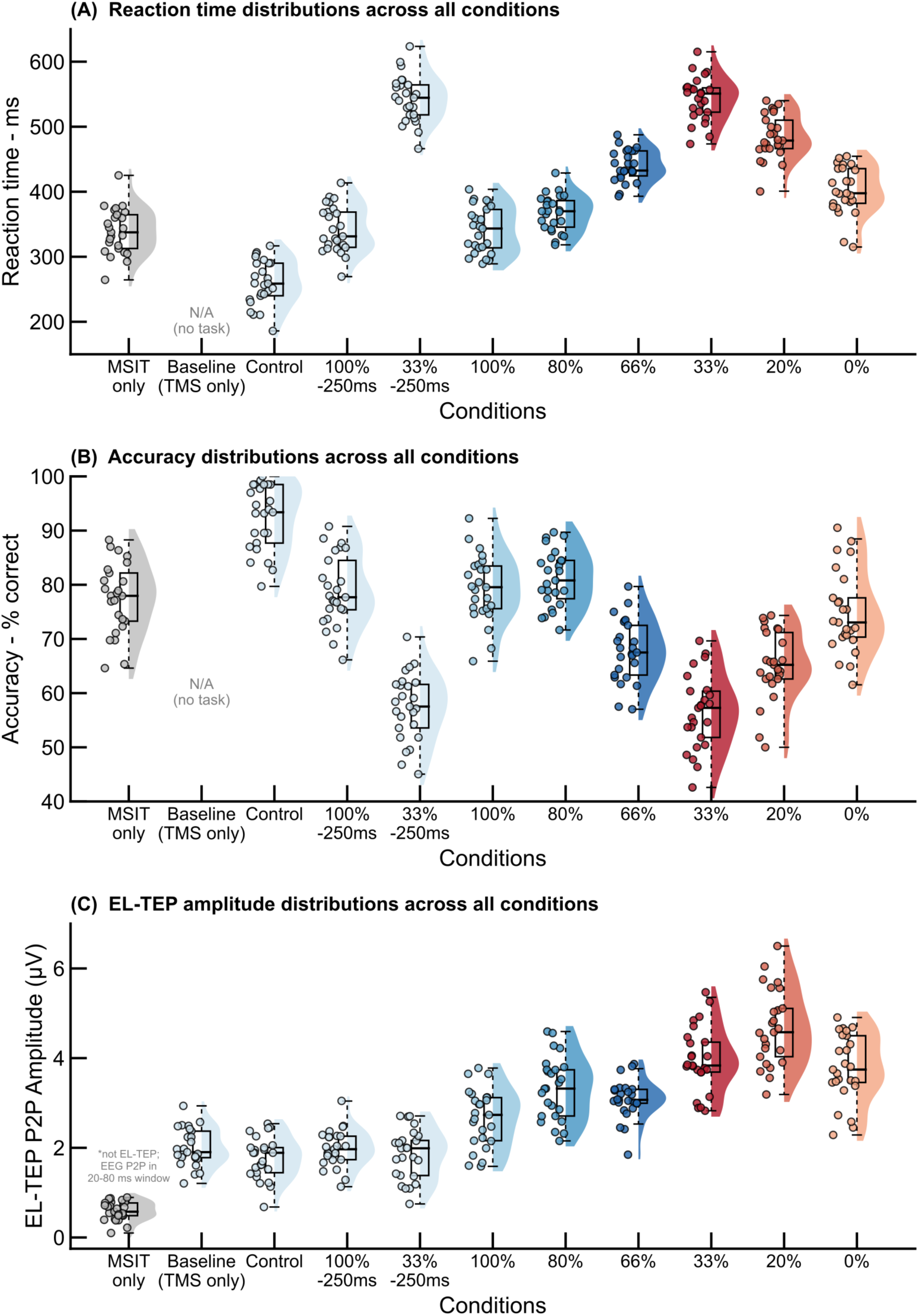
Individual subject behavioral (reaction times & accuracy) and EL-TEP distributions across all 11 experimental conditions of the main left-dlPFC TMS experiment. (A) Mean reaction time per subject for each condition (trials with RT < 750 ms). The Baseline condition (TMS-only) has no associated behavioral data. (B) Mean accuracy (% correct) per subject for each condition. (C) EL-TEP peak-to-peak amplitude (µV) per subject for each condition. Raincloud plots display kernel density estimates, box plots (median and interquartile range), and individual subject data points. Conditions are ordered as: ERP, Baseline, Control, 100% Control (-250 ms), 33% Control (-250 ms), 100%, 80%, 66%, 33%, 20%, and 0% congruent. N = 27 subjects. Data include only trials with reaction times <750 ms to focus on valid responses and exclude outliers.

**Figure S2.**
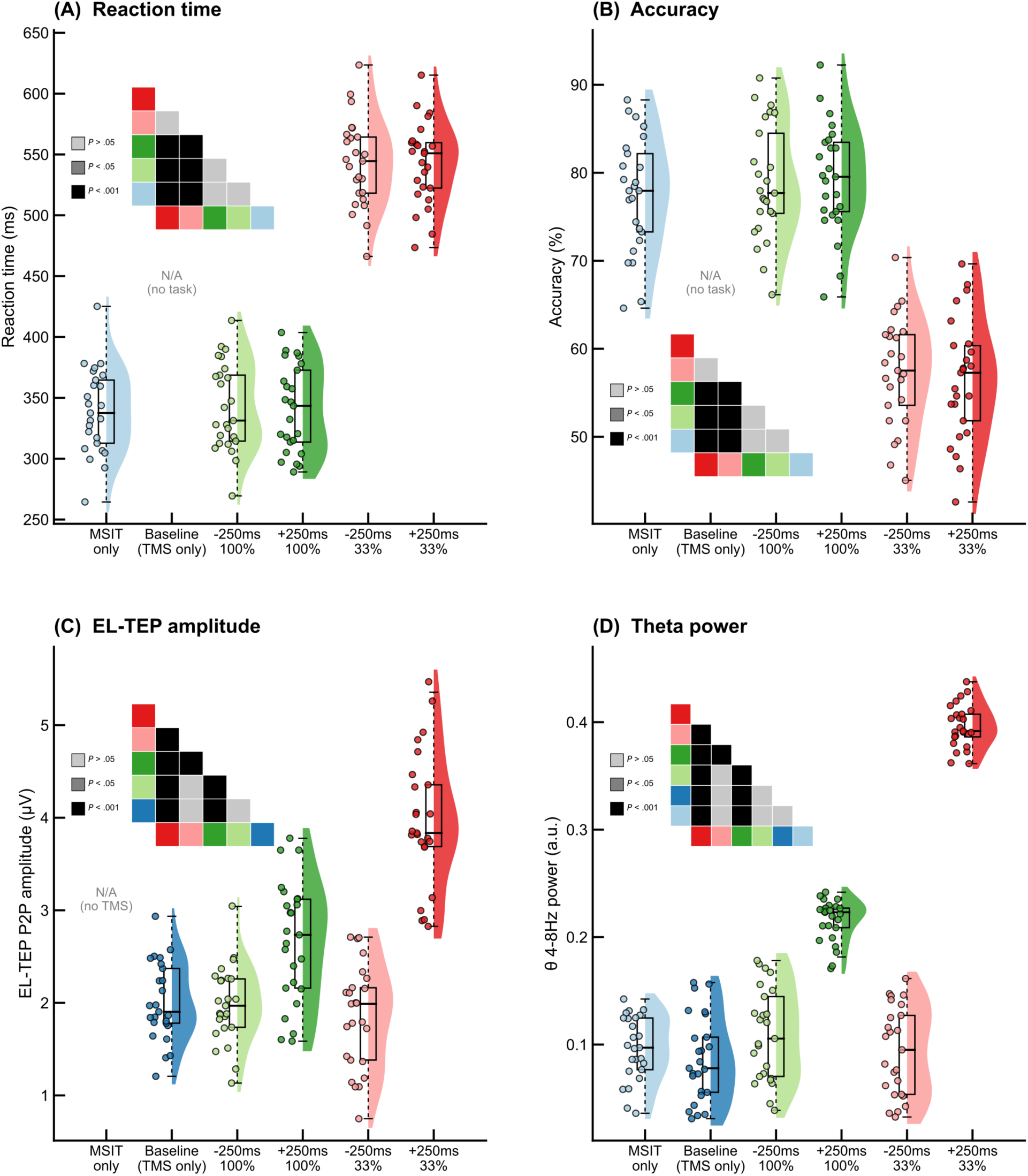
TMS timing effects on prefrontal excitability relative to MSIT stimulus onset. (A) Mean reaction time per subject for each timing condition. (B) Mean accuracy (% correct) per subject for each timing condition. (C) EL-TEP peak-to-peak amplitude (µV) per subject for each timing condition. (D) Theta (4–8 Hz) spectral power per subject for each timing condition. A selective set of conditions is shown to examine the effect of TMS timing relative to MSIT stimulus onset. The −250 ms conditions delivered TMS 250 ms before stimulus onset, while +250 ms conditions delivered TMS +250 ms after stimulus onset, bracketing the critical period of cognitive interference processing. Raincloud plots display kernel density estimates, box plots (median and interquartile range), and individual subject data points. Significance matrix (top left) indicates pairwise comparisons. N = 27 participants.

**Figure S3.**
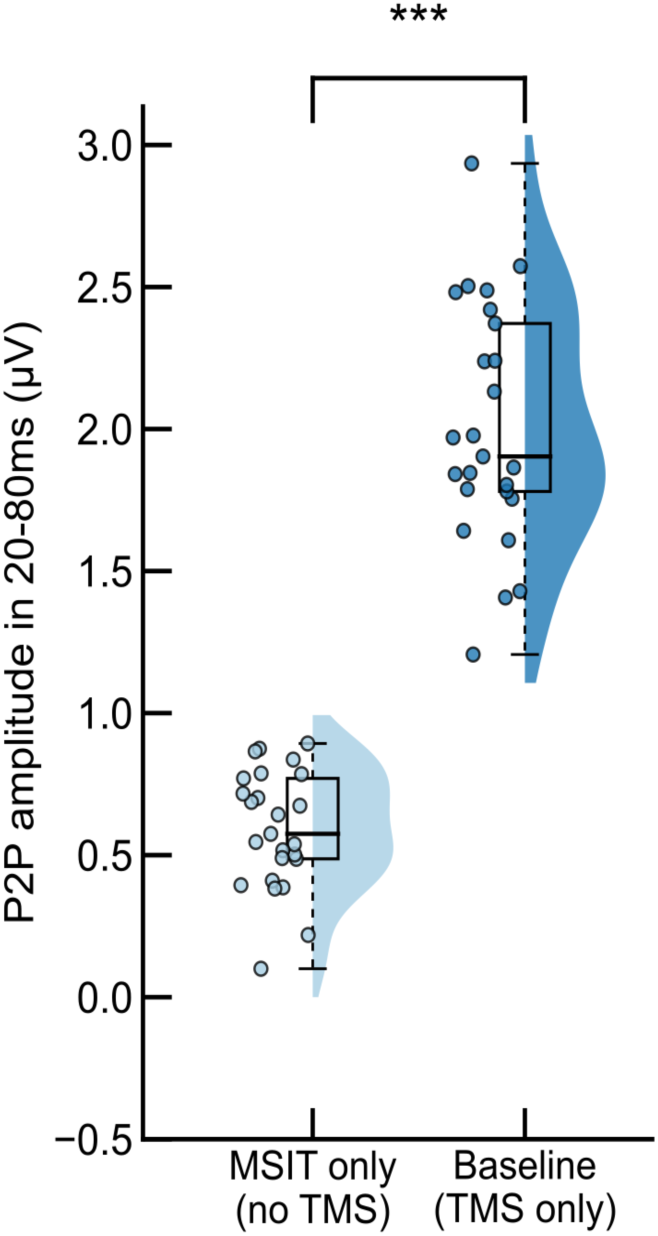
Comparison of spontaneous prefrontal EEG activity in MSIT only condition and TMS-evoked prefrontal response amplitude in TMS only condition. Spontaneous prefrontal EEG (no TMS) versus TMS-evoked prefrontal activity (EL-TEP) measured in the same 20–80 ms post-stimulation window. Raincloud plots display kernel density estimates, box plots (median and interquartile range), and individual subject data points. The MSIT only condition reflects task-related EEG activity without TMS delivery, while the TMS only (baseline) condition reflects the early-local TMS-evoked response (EL-TEP) amplitude in the absence of a cognitive task. ***P < 0.001.

**Figure S4.**
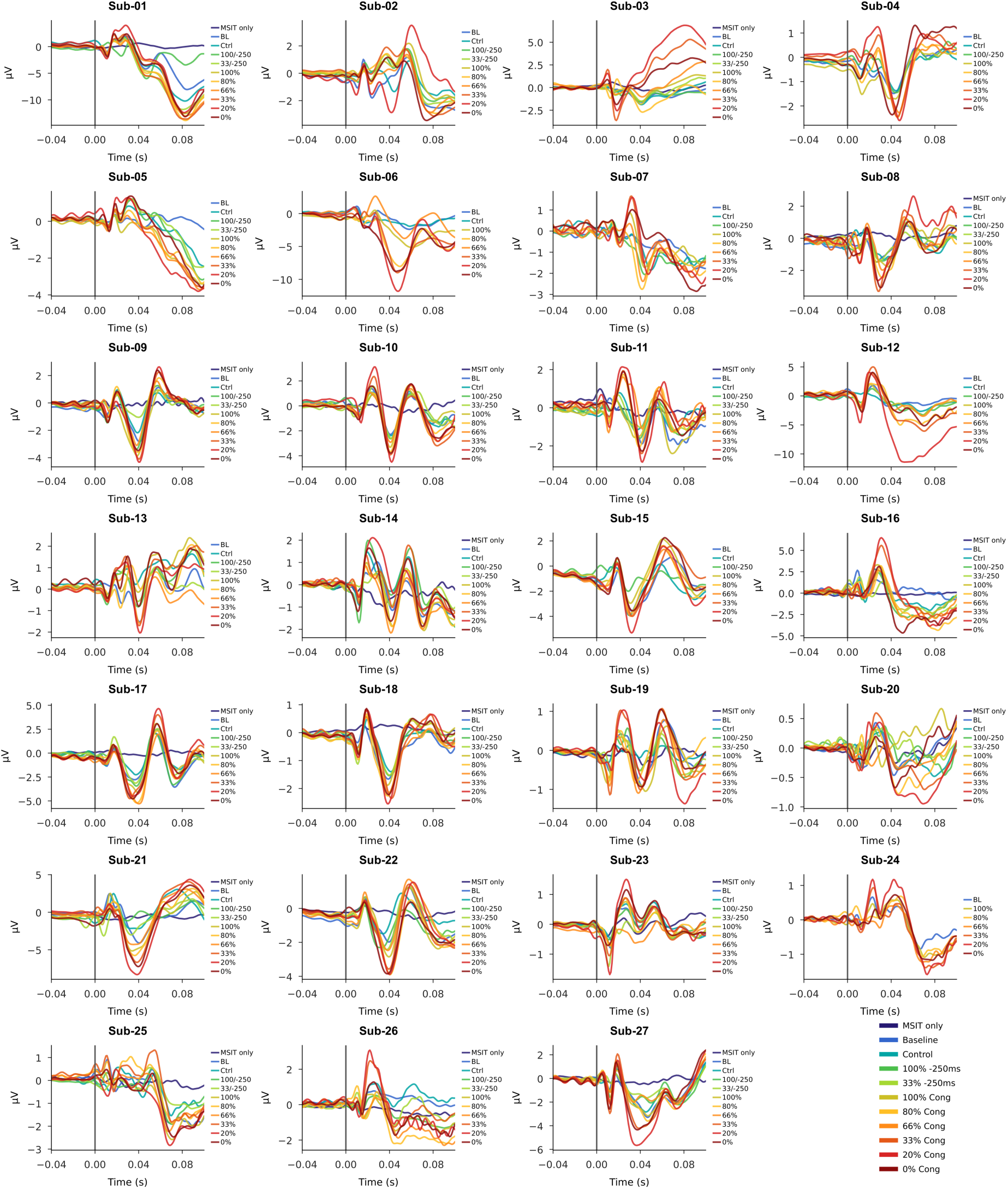
Modulation of EL-TEPs in left dLPFC reveals subject-level condition-specific neurophysiological signatures across cognitive interference. EL-TEPs recorded from the left dLPFC under various cognitive interference conditions from -0.05s to 0.10s relative to TMS pulse (dashed line at 0.00s). Different colored traces represent experimental conditions: baseline, control task, and varying levels of congruent and control trials as shown in the legend. Notable modulation of EL-TEP components occurs, particularly the negative deflection (0.03-0.04s) and positive peak (0.06-0.07s), demonstrating how TMS-evoked brain activity in the left dLPFC is differentially affected by cognitive load.

**Figure S5.**
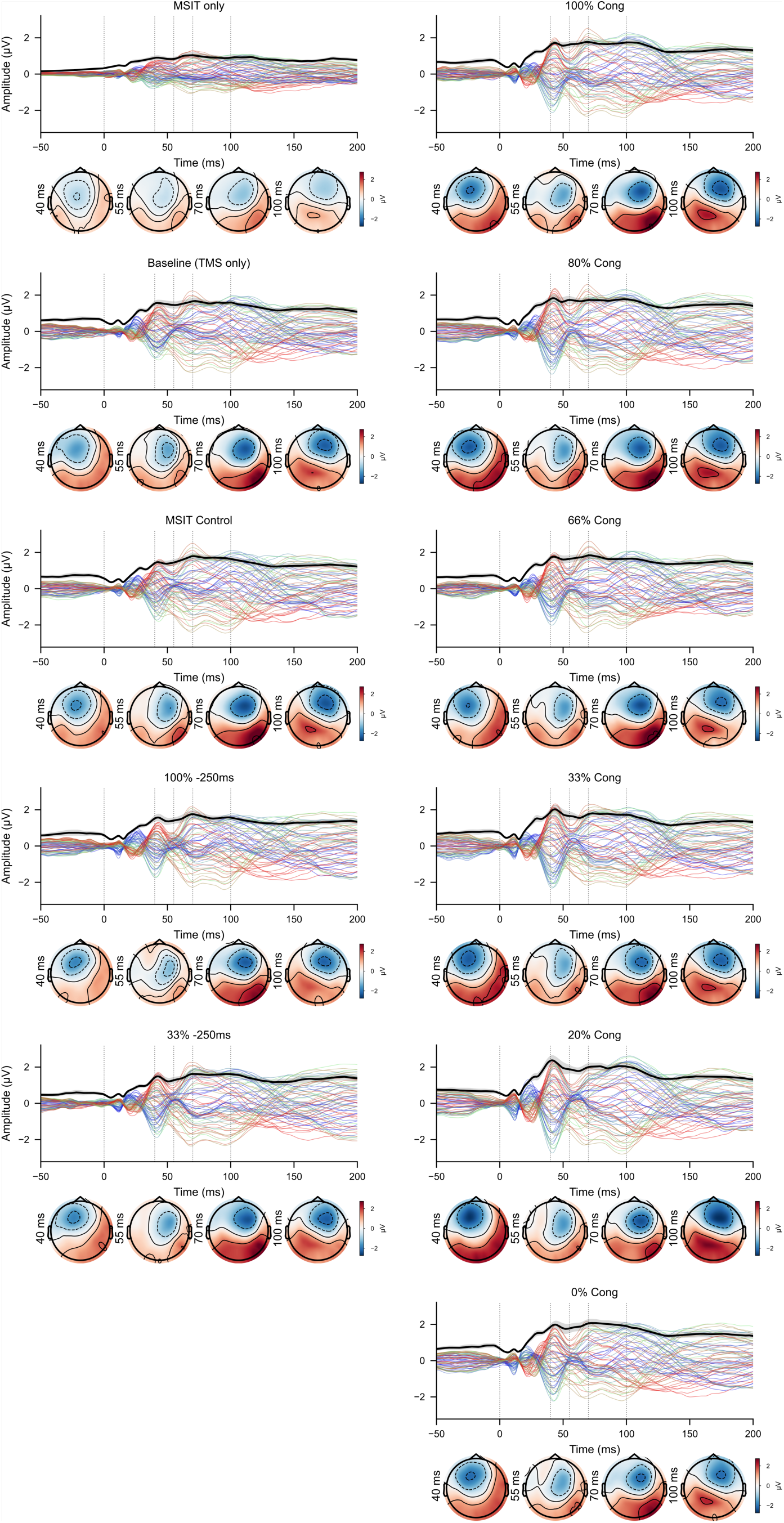
Grand-average butterfly plots and scalp topographies for all experimental conditions across all subjects. Each panel shows the grand-average TMS-evoked potential across all 63 EEG channels (colored traces) with the global mean field amplitude (GMFA; black line ± SEM shading). Scalp topographies are shown at 40, 55, 70, and 100 ms post-TMS pulse.

**Table S1.** Demographics. *N = 32. Participants were on average 40.21 years old (SD = 12.87)*.

| Characteristic | n | % of sample |
| --- | --- | --- |
| Gender |  |  |
| Female | 13 | 42.11 |
| Male | 18 | 55.26 |
| Nonbinary | 1 | 2.63 |
| Race |  |  |
| Asian | 10 | 31.58 |
| Black or African American | 4 | 13.16 |
| White | 15 | 44.74 |
| Other | 3 | 10.53 |
| Handedness |  |  |
| Left | 2 | 5.26 |
| Right | 30 | 94.74 |
| Highest education level |  |  |
| Completed high school | 1 | 2.63 |
| Some college | 3 | 10.53 |
| 2-year college | 4 | 13.16 |
| 4-year college | 12 | 36.84 |
| Pursuing post graduate degree | 3 | 7.89 |
| Post graduate degree | 9 | 28.95 |

**Table S2.** Stimulation parameters across participants (N = 32)

| **Participant ID**<br>**(Sub-XX)** | **rMT** | **Optimized angle** | **Highest Intensity**<br>**% of rMT (leftM1)** | **% MSO for**<br>**highest intensity** |
| --- | --- | --- | --- | --- |
| 02 | 48 | -100.7 | 140 | 67 |
| 03 | 53 | -11.3 | 140 | 74 |
| 04 | 56 | -42.4 | 140 | 78 |
| 06 | 52 | -79.2 | 140 | 73 |
| 07 | 70 | -73.1 | 140 | 98 |
| 08 | 61 | N/A | 140 | 85 |
| 09 | 63 | -58.9 | 140 | 88 |
| 10 | 82 | -30.8 | 121 | 100 |
| 11 | 70 | N/A | 140 | 98 |
| 12 | 69 | N/A | 140 | 97 |
| 13 | 57 | N/A | 140 | 80 |
| 14 | 70 | N/A | 140 | 98 |
| 15 | 54 | N/A | 140 | 76 |
| 16 | 64 | N/A | 131 | 84 |
| 17 | 50 | N/A | 140 | 70 |
| 18 | 77 | N/A | 119 | 92 |
| 19 | 73 | N/A | 136 | 100 |
| 20 | 66 | 11.8 | 140 | 92 |
| 21 | 72 | -11.1 | 138 | 100 |
| 22 | 60 | -14.4 | 140 | 84 |
| 23 | 62 | -64.5 | 135 | 84 |
| 24 | 55 | 22.8 | 140 | 77 |
| 25 | 55 | -97.5 | 140 | 77 |
| 26 | 60 | -63.3 | 140 | 84 |
| 27 | 85 | -77.2 | 117 | 100 |
| 28 | 49 | LDLPFC: -24.7<br>RDL PFC: N/A<br>LPreSMA: -100<br>Vertex: 0 | 140 | 69 |
| 29 | 69 | LDLPFC: -54.3<br>RDL PFC: 78.2<br>LPreSMA: 46.2<br>Vertex:3.07 | 140 | 97 |
| 30 | 49 | LDLPFC N/A<br>RDL PFC: 8.06<br>LPreSMA: -100<br>Vertex:0 | 140 | 69 |
| 31 | 72 | LDLPFC: -22.3<br>RDLPFC: 32.9<br>LPreSMA: -100<br>Vertex: 0 | 138 | 100 |
| 32 | 71 | LDLPFC: -41.1<br>RDLPFC: N/A<br>LPreSMA: -100<br>Vertex: 0 | 140 | 100 |

## Notes

**Funding:** This work was supported by R01MH129018, R01MH126639, and the Burroughs Wellcome Fund Career Award for Medical Scientists.

### Competing Interest Statement

C.J. Keller holds equity in Alto Neuroscience, Inc. and Flow Neuroscience, Inc. All other authors declare no competing interests.

